# Brown adipose expansion and remission of glycemic dysfunction in obese SM/J mice

**DOI:** 10.1101/724369

**Authors:** Caryn Carson, Juan F Macias-Velasco, Subhadra Gunawardana, Mario A Miranda, Sakura Oyama, Heather Schmidt, Jessica P Wayhart, Heather A Lawson

**Affiliations:** Department of Genetics, Washington University School of Medicine, 660 South Euclid Ave, Saint Louis, MO, USA; Department of Cell Biology and Physiology, Washington University School of Medicine, 660 South Euclid Ave, Saint Louis, MO, USA; Department of Medicine, Washington University School of Medicine, 660 South Euclid Ave, Saint Louis, MO, USA

**Keywords:** glycemic control, brown adipose, obesity, metabolism, mouse model, transcriptome

## Abstract

Disruption of glucose homeostasis increases the risk of type II diabetes, cardiovascular disease, stroke, and cancer. We leverage a novel rodent model, the SM/J mouse, to understand glycemic control in obesity. On a high fat diet, obese SM/J mice initially develop impaired glucose tolerance and elevated fasting glucose. Strikingly, their glycemic dysfunction resolves by 30 weeks of age despite persistence of obesity. A prominent phenotype is that they dramatically expand their brown adipose depots as they resolve glycemic dysfunction. This occurs naturally and spontaneously on a high fat diet, with no temperature or genetic manipulation. When the brown adipose depot is removed from normoglycemic obese mice, fasting blood glucose and glucose tolerance revert to unhealthy levels, and animals become insulin resistant. We identified 267 genes whose expression changes in the brown adipose when the mice resolve their unhealthy glycemic parameters, and find the expanded tissue has a ‘healthier’ expression profile of cytokines and extracellular matrix genes. We describe morphological, physiological, and transcriptomic changes that occur during the unique brown adipose expansion and remission of glycemic dysfunction in obese SM/J mice. Understanding this phenomenon in mice will open the door for innovative therapies aimed at improving glycemic control in obesity.

**Significance Statement:** Some obese individuals maintain normal glycemic control. Despite being obese, these individuals have low risk for metabolic complications, including type-II diabetes. If we better understood why some obese people maintain normoglycemia then we might develop new approaches for treating metabolic complications associated with obesity. However, the causative factors underlying glycemic control in obesity remain unknown. We discovered that, despite persistence of the obese state, SM/J mice enter into diabetic remission: returning to normoglycemia and reestablishing glucose tolerance and improving insulin sensitivity. A prominent phenotype is that they dramatically expand their brown adipose depots as they resolve glycemic dysfunction. Understanding this phenomenon in mice will open the door for innovative therapies aimed at improving glycemic control in obesity.

## Introduction

An estimated 10-30% of obese individuals maintain glycemic control and some longitudinal studies suggest their risk of developing type II diabetes is no greater than matched lean individuals (1). No causative factors underlying glycemic control in obesity have been discovered, however the strongest predictors of impaired glycemic control in obesity are increased visceral fat mass and adipose tissue dysfunction (2,3). Thus research efforts have focused on understanding the genetic and physiological mechanisms of action of adipose. Recent research reveals that brown adipose activity is associated with anti-diabetic properties. Cold exposure in both obese and lean individuals causes increased uptake of fatty acids and glucose into brown adipose tissue (4). Further, increased brown adipose activity has been shown to improve glucose homeostasis and insulin sensitivity in adults (5). Transplantation of brown adipose tissue into mouse models of diabetes greatly improves glucose parameters, including fasting glucose levels and response to a glucose challenge (6). While there are a variety of obese and diabetic mouse models, there are no mouse models for understanding the relationship between brown adipose and glycemic control in obesity.

The SM/J inbred mouse strain has long been used for studying interactions between diet and metabolism, and more recently has started to help uncover the genetic architecture underlying diet induced obesity and glucose homeostasis. It has previously been shown that fed a high fat diet, SM/J mice display many of the characteristics of a diabetic-obese mouse: obesity, hyperglycemia and glucose intolerance at 20 weeks of age (7,8). We discovered that SM/J mice undergo a remarkable transformation between 20 and 30 weeks of age. Despite persistence of the obese state, these mice enter into diabetic remission: returning to normoglycemia and reestablishing glucose tolerance and improving insulin sensitivity. Contemporary with this remission of glycemic parameters is a dramatic expansion of the intrascapular brown adipose depot. This study describes the morphological, physiological, and transcriptomic changes that occur during this transition, and establishes the SM/J mouse as a unique model for understanding the relationship between brown adipose and glycemic control in obesity. Understanding this relationship in a genetic model of glycemic resolution will set the stage for identifying novel, potentially therapeutic targets for the improvement of glycemic control.

## Results

### SM/J mice improve glucose parameters without weight loss

When fed a high fat diet (**Supplemental Table 1**) from 3 weeks of age, SM/J mice develop obesity, hyperglycemia, and impaired glucose tolerance by 20 weeks (9). By 30 weeks, despite the persistence of obesity, high fat-fed SM/J’s resolve their hyperglycemia and impaired glucose tolerance to levels indistinguishable from low fat-fed controls (**Figure 1A-D**). Further, 30 week old high fat-fed SM/J mice improve insulin sensitivity (**Figure 1E and F**).

**Figure 1.**
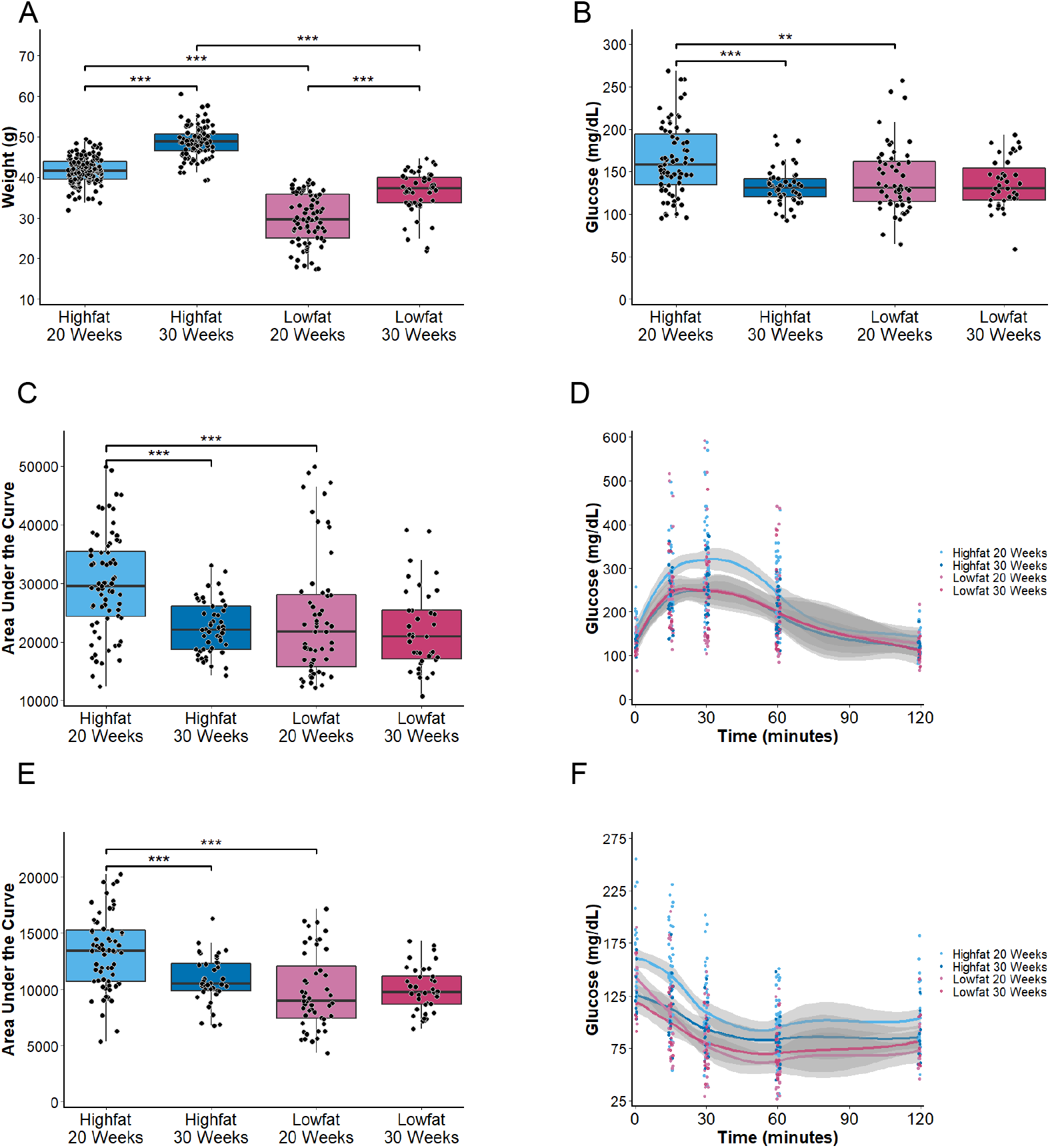
Obese SM/J mice improve glucose parameters between 20 and 30 weeks of age. **A** High fat-fed mice weigh significantly more than low fat-fed mice, and SM/J mice gain weight between 20 and 30 weeks of age on both diets, n = 48-131 mice per cohort. **B** 30 week-old high fat-fed mice have significantly lower fasting glucose levels than at 20 weeks, which is no different than low fat-fed controls, n = 22-47 mice per cohort. **C and D** 30 week-old high fat-fed mice have improved glucose tolerance relative to 20 weeks, n = 22-47 mice per cohort. **E and F** Lower fasting glucose and improved glucose tolerance corresponds with improved insulin sensitivity, n = 22-47 animals per cohort. Equal numbers of males and females represented; * p<0.05, ** p<0.01, *** p<0.001

High fat-fed C57BL/6J mice also show a reduction in fasting glucose that is accompanied by increased insulin with age (10). In contrast to SM/J, the difference in circulating glucose between the high fat- and low fat-fed C57BL/6J remain significantly different over time. Moreover, high fat-fed C57BL/6J mice show marked glucose intolerance that does not resolve with age. We observe a similar trend in the LG/J strain of mice, where high fat-fed animals maintain higher fasting glucose levels and impaired glucose tolerance relative to low fat-fed controls as they age (**Supplemental Figure 1**). The unique remission of hyperglycemia and improved glucose tolerance observed in the high fat-fed SM/J strain indicates a genetic basis.

### High fat-fed SM/J mice expand their interscapular brown adipose tissue depots

Contemporary with the resolution of glycemic parameters, high fat-fed SM/J mice dramatically expand their intrascapular brown adipose depots, which is not seen in low fat-fed control mice (**Figure 2A-C**). This has never been described in another mouse strain, and we do not observe the phenomenon in the LG/J strain of mice on the same diets at any age (**Supplemental Figure 2**). To understand whether the tissue mass expansion is due to increased size of individual cells or to increased number of total cells, we quantified adipocyte cell size and the mitotic index. There are no significant differences in average cell size in high fat-fed mice between 20 and 30 weeks, or relative to low fat-fed controls (**Supplemental Figure 3A**). Mice on both diets undergo altered adipocyte area profiles between 20 and 30 weeks of age, however the low fat tissue develops a profile significantly trending towards larger adipocytes at 30 weeks (p=6.4^-07^) whereas the high fat tissue develops a profile significantly trending towards smaller adipocytes at 30 weeks (p=2.2^-16^) (**Figure 2D and E**). This suggests that the expansion of the brown adipose depot in high fat-fed mice is the result of increased proliferation of adipocytes, as newer adipocytes are smaller due to less lipid accumulation. This is supported by quantification of brown adipose cells stained positive for the mitotic marker phosphohistone H3, which trends towards a higher mitotic index in the brown adipose of high fat-fed animals (**Supplemental Figure 3B**).

**Figure 2.**
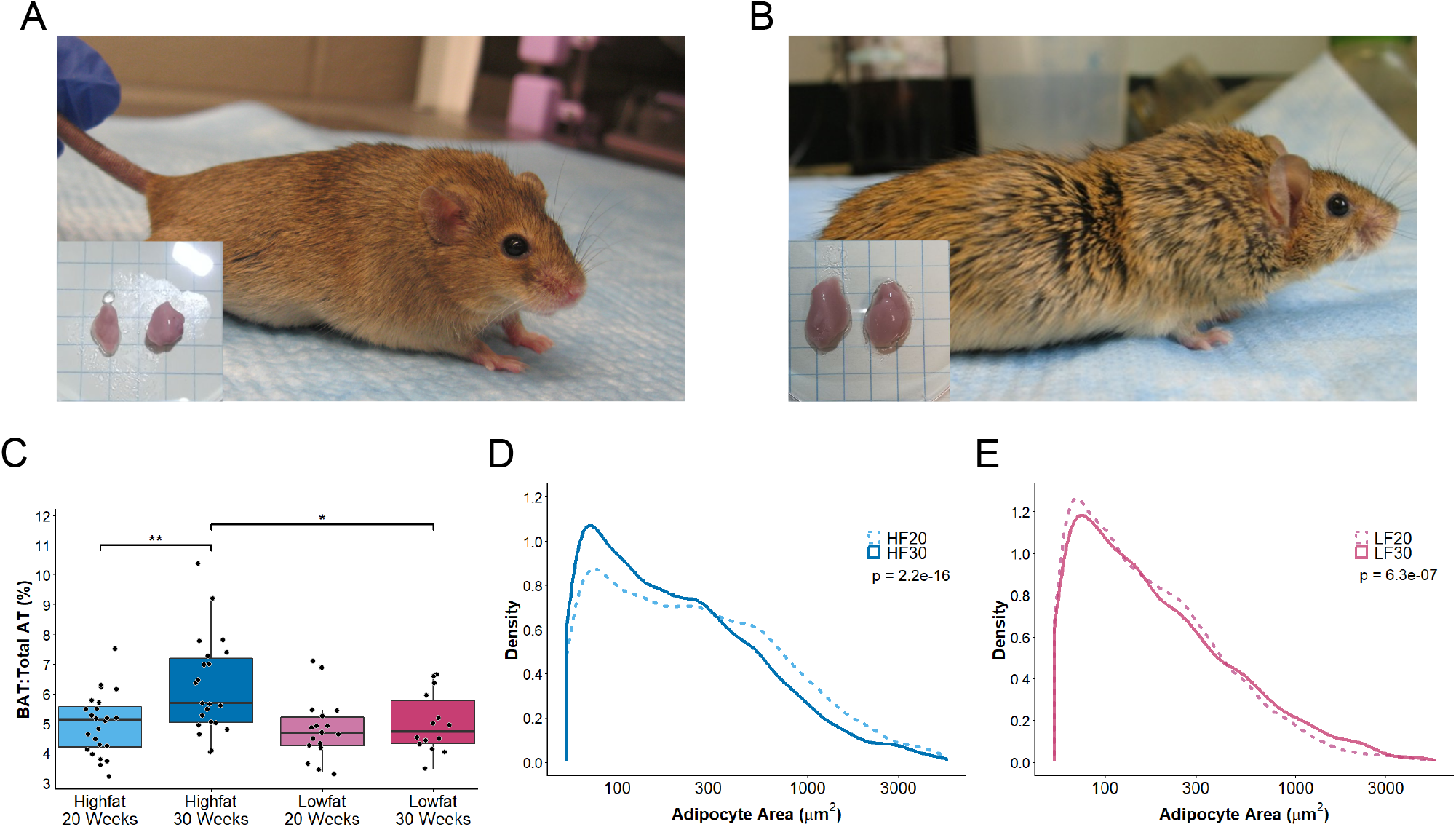
Brown adipose expansion in 30 week-old high fat-fed SM/J mice. Representative pictures of **A** 20 week and **B** 30 week-old high fat-fed female mice. **C** Quantification of interscapular brown adipose depot as a proportion of total fat mass, n = 16-25 mice per cohort. **D and E** Cell area density graphs for high fat and low fat-fed cohorts. Data are plotted on a log 10 scale for visualization, n = 4 mice per cohort.

Because obesity has been associated with structural and functional “whitening” of brown adipose depots in rodents (11–14), we performed experiments to confirm that the tissue expansion in SM/J mice has the expected properties of brown fat. Histological analysis of the fat depot taken from high fat-fed SM/J mice at 30 weeks of age confirms the adipocytes in this expansion are brown adipocytes, with small multilocular lipid droplets and high UCP1 staining (**Supplemental Figure 3C-J**). Expression of canonical brown adipose genes *Ucp1* and *Cidea* do not change between 20 and 30 weeks (**Supplementary Figure 4A-B**). Further, expression of *Tbx1*, a marker specific for beige adipocytes (15), indicates that neither brown nor white adipose is “beiging” (**Supplementary Figure 4C**). Finally, there is no significant difference in brown adipose tissue mitochondrial content between the diets or ages (**Supplementary Figure 4D**). There is no difference in core body temperature or circulating free fatty acids between high and low fat-fed cohorts or between 20 and 30 weeks of age (**Supplemental Figure 5A and B**). Additionally, while there are diet-dependent differences in the catecholamines norepinephrine and epinephrine, which activate UCP1-mediated leak respiration and non-shivering thermogenesis, there is no change in levels between ages in the high fat-fed mice (**Supplemental Figure 5C and D**). Thus, the interscapular adipose depot in high fat-fed SM/J mice maintains a brown adipose identity after expansion that is not dependent on whole-animal beiging, and is also not associated with altered thermogenesis.

### Glucose parameters revert to an unhealthy state in SM/J mice when the brown adipose depot is removed

If the brown adipose expansion is directly related to the glycemic resolution of the high fat-fed SM/J mice, preventing or removing that expansion should revert the glucose parameters to their unhealthy state. To test these predictions, we removed the interscapular brown adipose depots from hyperglycemic 20 and normoglycemic 30 week-old mice. After recovery, at 30 and 35 weeks of age, we measured basal glucose levels and performed glucose and insulin tolerance tests. We find that improvement in glucose and insulin tolerance is prevented when the brown adipose depot is removed before expansion. Previously normoglycemic mice revert to unhealthy, 20 week-old measurements when the brown adipose depot is removed after expansion (**Figure 3 A-C**). These results indicate that the expanded brown adipose tissue is necessary for the observed remission of glucose intolerance, and for the maintenance of both glucose tolerance and insulin sensitivity in high fat-fed SM/J mice.

**Figure 3.**
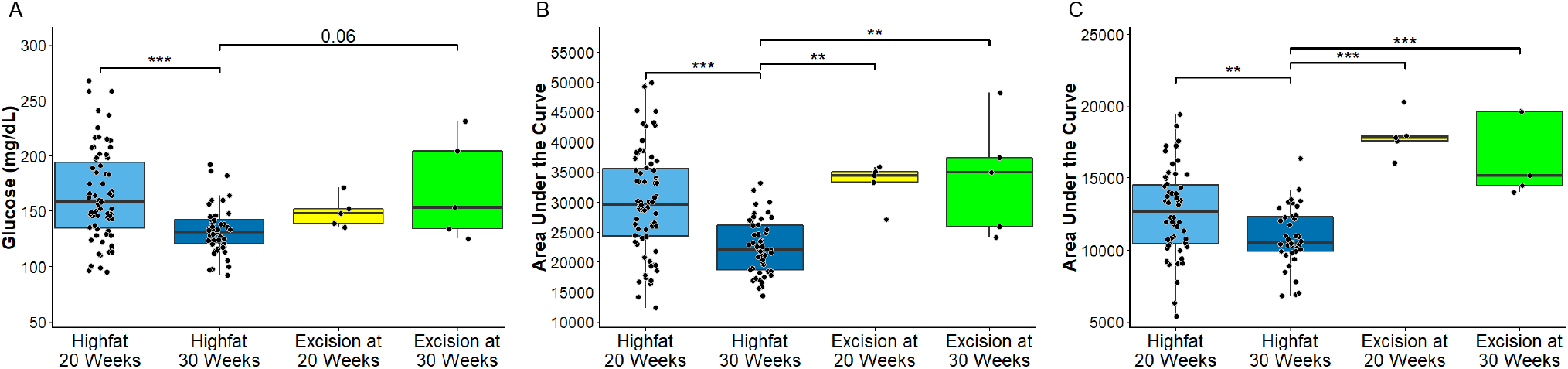
Glycemic parameters revert to unhealthy levels when the brown adipose depot is removed. **A** Blood glucose assessed after a 4 hour fast. **B** A glucose tolerance test indicates removal of the brown adipose depot before (20 week excision) or after (30 week excision) expansion significantly reduces glucose tolerance. **C** Removal of the brown adipose tissue also significantly increases insulin resistance. n = 5 excision animals per cohort, representing 4 males and 1 female.

### RNA sequencing reveals enrichment of differentially expressed cytokines and genes affecting extra cellular matrix

Since the brown adipose tissue expansion is unique to high fat-fed SM/J mice, we anticipated that there would be corresponding unique transcriptomic changes in the brown adipose. Indeed, we identified 267 genes whose expression significantly and uniquely changes between 20 and 30 weeks of age in high fat-fed SM/J brown adipose tissue (at a 5% FDR, out of 13,253 total genes expressed; **Supplemental Table 2**). These expression changes occur when the mice resolve their glycemic dysfunction and expand their brown adipose depots. These genes are not differentially expressed in white adipose tissue taken from the same animals or in low fat-fed SM/J controls (**Figure 4A**). Additionally, they are not differentially expressed in the LG/J strain of mouse, once again underscoring the genetic basis of the phenomenon (**Supplemental Table 3**).

**Figure 4.**
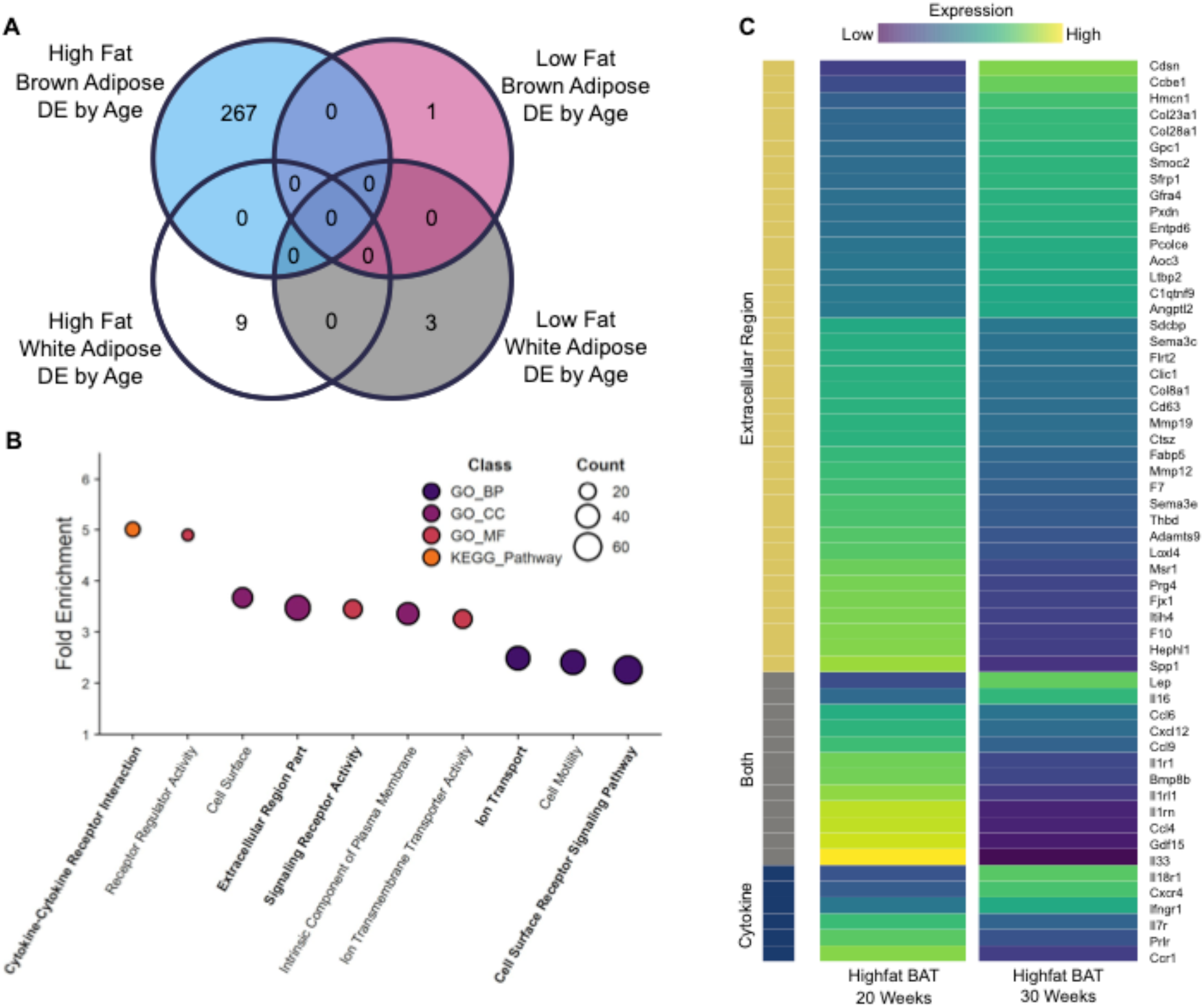
High fat-fed SM/J mice have unique brown adipose differential expression between 20 and 30 weeks of age. **A** Venn diagram illustrating the number of genes differentially expressed between high and low fat-fed 20 and 30 week-old SM/J interscapular brown or reproductive white adipose tissues. No genes are differentially expressed in more than one diet-by-tissue cohort. **B** Enriched terms colored by class (Gene Ontology Biological Process (GO_BP), Cellular Component (GO_CC), Molecular Function (GO_MF), and KEGG Pathway. **C** Heatmap of differentially expressed brown adipose tissue genes between high fat-fed 20 and 30 week-old mice belonging to cytokine, extracellular matrix, or both gene ontologies. Equal numbers of males and females represented, n = 8 animals per age-by-diet cohort.

Over-representation analysis indicates these genes are enriched for those involved in cytokine-cytokine receptor interactions (p=3.23e^-06^), signaling receptor activity (p = 5.70e^-06^), cell surface receptor signaling (p=2.04e^-07^), and extracellular matrix components (p = 7.93 e^-13^) (**Figure 4B**). These are intriguing results because brown adipose has been identified as a source of cytokines that influence glucose homeostasis, and extracellular matrix changes are essential for tissue expansion, cellular signaling, and regulation of growth factor bioavailability.

Several genes belonging to these biological categories have evidence for their involvement in glucose homeostasis and change expression in a direction that is associated with improved metabolic health in high fat-fed SM/J mice between 20 and 30 weeks of age (**Figure 4C; Supplemental Table 2**). In particular, the direction of expression change reveals that the expansion of brown adipose is associated with decreased expression of inflammatory (e.g. interleukin 7 receptor, *Il7r*) (16) and fibrotic markers (e.g. collagen type VIII alpha 1 chain, *Col8a1;* semaphorin 3C, *Sema3c*) (17,18), and changes in extracellular matrix components (e.g. matrix metallopeptidase 12, *Mmp12;* procollagen c-endopeptidase enhancer, *Pcolce*) (19,20) and cytokines (e.g. coagulation factor VII, *F7;* leptin, *Lep;* secreted frizzled-related protein 1, *Sfrp1*) (21–23). These genes are not differentially expressed in white adipose from the same animals, and they are not differentially expressed in low fat-fed brown or white adipose between 20 and 30 weeks (**Supplemental Figure 6**). Other mouse models of diet-induced obesity develop unhealthy brown adipose transcriptomes characterized by increased expression of pro-inflammatory genes and fibrotic markers (24). The direction of expression change in our brown adipose tissue supports the uniqueness of the high fat-fed SM/J mice.

### 30 week old high fat-fed SM/J brown adipose has a healthier co-expression profile

Because variation in glucose homeostasis is complex and the result of many interacting genes, we examined the co-expression profile of genes belonging to the enriched cytokine and extracellular matrix (ECM) biological categories (**Figure 4B and C**). We find that the co-expression profile of the differentially expressed ECM and cytokine genes is significantly different between 20 and 30 week-old high fat-fed animals (p=0.01). To determine if the co-expression profile of these genes in 30 week-old high fat-fed animals’ brown adipose is more similar to the 20 week-old high fat-fed or to the low fat-fed animals’, we compared the overall co-expression correlation structure between the diet and age cohorts for these genes. Remarkably, we find the 30 week-old high fat-fed SM/J brown adipose ECM and cytokine co-expression profile is most similar to the 20 week-old low fat-fed animals’ (probability of difference between high fat-fed 30 weeks and low fat-fed 20 weeks = 0.07; probability of difference between high fat-fed 30 weeks and low fat-fed 30 weeks = 0.04) (**Figure 5A**). Thus, the brown adipose cytokine and ECM gene co-expression profile appears ‘healthier’ in 30 week-old high fat-fed animals after expansion and remission of the diabetic phenotype. This is illustrated in **Figure 5B**.

**Figure 5.**
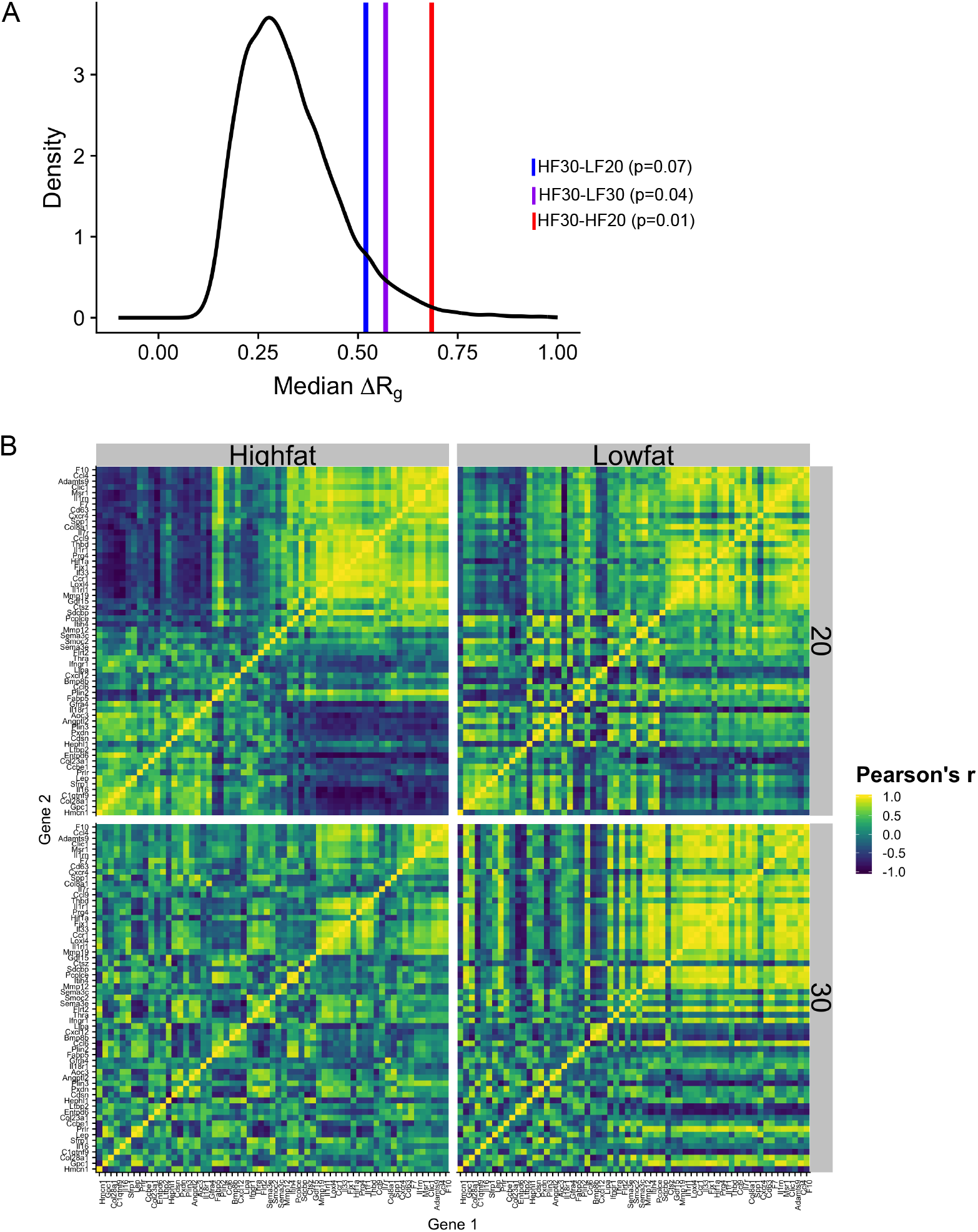
High fat-fed 30 week-old SM/J mice have a cytokine and ECM gene co-expression profile most similar to low fat-fed 20 week-old mice. **A** The median change in correlation structure is plotted as a vertical line against the null model. High fat-fed 30 week-old SM/J mice have a co-expression profile significantly different from high fat-fed 20 week-old SM/J mice, and not significantly different from low fat-fed 20 week-old mice. **B** Heatmap of the gene expression correlation matrices for each cohort. HF30 = high fat-fed 30 week-old; HF20 = high fat-fed 20 week-old; LF30 = low fat-fed 30 week-old; LF20 = low fat-fed 20 week-old. Equal numbers of males and females represented, n = 8 animals per age-by-diet cohort.

## Discussion

Obesity (body-mass index [BMI] ≥30 kg.m^2^) is associated with serious metabolic complications, including type II diabetes, cardiovascular disease, cancer, and stroke (25–29). Currently, 38% of American adults are classified as obese, 9.4% have type II diabetes, and an additional 34% are prediabetic, costing 327 billion dollars in annual medical costs (30,31). Obesity and diabetes are tightly linked; obesity raises the risk of developing type II diabetes 27–76 fold, while approximately 60% of diabetics are obese (25,30,32,33). Though weight loss is *the* gold standard for treating glycemic dysfunction in obesity, many obese people are unable to achieve long-term weight loss (34,35). Currently, metformin and thiazolidinediones are prescribed to prevent the development of diabetes in obese individuals, but pharmacological therapy has been shown to have only modest protective effects (33,36). Greater understanding of the relationship between obesity and glycemic control is needed to develop more effective preventative measures for obese patients.

Here we describe morphological, physiological, and transcriptomic changes that occur during brown adipose expansion and remission of glycemic dysfunction in obese SM/J mice. The SM/J strain was derived from a pool of seven inbred strains and selected for small body size at 60 days (37). The strain has been used extensively in genetic studies of complex traits related to growth and metabolism, particularly because SM/J mice are strongly responsive to high fat diet-induced obesity. These studies all used mice aged 20 weeks or less, when SM/J mice develop the classic hallmarks of obese-diabetic mice. We discovered that by 30 weeks of age, contemporary with a dramatic expansion of intrascapular brown adipose tissue, their hyperglycemia and impaired glucose tolerance go into remission despite persistence of obesity. Dissecting the genetic basis of this phenomenon has the potential to uncover novel relationships among brown adipose, glucose homeostasis, and obesity.

We identified 267 genes whose expression significantly and uniquely changes between 20 and 30 weeks of age in high fat-fed SM/J brown adipose tissue (**Figure 5A**). We focus on genes associated with ECM and cytokine activity because both biological categories are enriched in the set of genes that significantly change expression in brown adipose during the remission of glycemic parameters. Brown adipose is a source of endocrine signals with anti-diabetic properties and is involved in extensive crosstalk with other organs (38). It secretes cytokines that influence whole-body glucose homeostasis and insulin sensitivity including IGF1, FGF21, NRG-3 and NRG-4 (39). ECM changes are essential for cellular signaling, regulation of growth factor bioavailability, and accompany healthy adipose expansion. However, extreme changes in ECM protein levels are associated with adipose dysfunction in obesity; thus a fine balance between tissue remodeling and excessive accumulation of ECM proteins must be achieved to maintain adipose tissue homeostasis (18).

We highlight 8 cytokines and ECM genes that significantly change expression in a direction associated with improved metabolic health in previous studies. *Il7r,* which was found to be one of the highest ranking genes in the white adipose tissue inflammatory response pathway (40), decreases expression between 20 and 30 weeks of age in high fat-fed SM/J brown adipose. *Col8a1* and *Sema3C* are both associated with adipose tissue fibrosis (17,41). Increased adipose tissue fibrosis is a signature of dysfunctional adipose and is associated with impaired glucose homeostasis and insulin resistance (18). Both *Col8a1* and *Sema3c* expression decrease between 20 and 30 weeks in high fat-fed SM/J brown adipose. *Mmp12* is an enzyme that contributes to adipose tissue remodeling (42). Increased *Mmp12* expression is associated with white adipose tissue inflammation and insulin resistance and *Mmp12^-/-^* mice are more insulin sensitive than wildtype controls on a high fat diet (19). Its expression decreases in 30 week old high fat-fed SM/J brown adipose. *Pcolce* encodes a glycoprotein that regulates collagen processing at the ECM (43). Mice with defects in ECM collagen are glucose intolerant, hyperglycemic, and insulin resistant (20). PCOLCE is one of 15 key drivers that collectively account for 22% of GWAS hits for type II diabetes in a recent multiethnic meta-analysis (44). *Pcolce* expression is significantly increased in 30 week old high fat-fed SM/J brown adipose. *F7, Lep,* and *Sfrp1* are each secreted proteins. Increased *F7* plays a role in the pathogenesis of obesity (45). In particular it has been shown to induce beta cell death and impaired islet glucose-stimulated insulin secretion (21). Increased *Lep* can dramatically lower blood glucose levels in diabetic rodent models (23). In brown adipose, leptin has been shown to stimulate glucose uptake (46). *Sfrp1* is dysregulated in obesity and *Sfrp1^-/-^* mice have elevated blood glucose and impaired glucose tolerance when fed a high fat diet (22). *F7* expression is decreased and *Lep* and *Sfrp1* are increased in 30 week old high fat-fed SM/J brown adipose tissue. Most of what is known about the role of these 8 genes in adipose comes from studies of white adipose tissue, but none of these genes are differentially expressed in SM/J white adipose. Many additional genes likely contribute to the observed phenomenon, however little, if anything, is known about their role in brown adipose tissue. The 267 differentially expressed genes we identified represent a set of actionable candidates for further functional studies of their role in brown adipose and glucose homeostasis.

There is great interest in harnessing the potential of brown adipose to treat obesity and diabetes, either through the calorie burning action of non-shivering thermogenesis or the endocrine action of adipokines. Research into the effects of brown adipose on systemic metabolism is in its infancy, and the community needs appropriate animal models to interrogate its physiological roles and identify potentially druggable targets. We present the SM/J mouse strain as a unique model to address this need. The SM/J mouse provides a tractable, genetic system in which to understand the relationship between brown adipose and glycemic control in obesity. Understanding this relationship in the SM/J mouse will open doors for identifying novel, potentially druggable targets for the improvement of glycemic control in humans.

## Methods

### Animal Husbandry and Phenotyping

SM/J (RRID:IMSR_JAX:000687) and LG/J (RRID:IMSR_JAX:000675) mice were obtained from The Jackson Laboratory (Bar Harbor, ME). Experimental animals were generated at the Washington University School of Medicine and all experiments were approved by the Institutional Animal Care and Use Committee in accordance with the National Institutes of Health guidelines for the care and use of laboratory animals. Pups were weaned at 3 weeks and reared in same-sex cages of 3-5 animals until necropsy. At weaning, mice were randomly placed on a high fat diet (42% kcal from fat; Teklad TD88137) or an isocaloric low fat diet (15% kcal from fat; Research Diets D12284) (**Supplemental Table 1**). Feeding was *ad libitum.* The animal facility operates on a 12 hour light/dark cycle with a constant ambient temperature of 21°C. Animals were weighed weekly until sacrifice. At 18 and 28 weeks of age, animals were subject to an intraperitoneal glucose tolerance test after a 4 hour fast. At 19 and 29 weeks of age animals were subject to an intraperitoneal insulin tolerance test. At 20 or 30 weeks of age, body composition was determined by MRI and temperature was measured with a rectal thermometer. After a 4 hour fast, at 20 or 30 weeks of age, animals were given an overdose of sodium pentobarbital and blood was collected via cardiac puncture. Euthanasia was achieved by cardiac perfusion with phosphate-buffered saline. After cardiac perfusion, tissues were collected and flash frozen in liquid nitrogen and stored at −80°C, or processed according to protocols for histology and other assays.

### Blood plasma assays

Fasting blood glucose was measured using a GLUCOCARD Vital glucometer (Arkay, MN USA). ELISAs measuring plasma levels of free fatty acids (Wako Life Sciences 995-34693) were quantified according to manufacturer’s protocol. Catecholamines were assayed through the Vanderbilt University Medical Center’s Hormone Assay and Analytical Services Core (www.vumc.org/hormone/assays; NIH grants DK059637 (MMPC) and DK020593 (DRTC)).

### Brown adipose histology

At the time of tissue collection, small portions of interscapular brown and reproductive white adipose tissues were placed in 1 mL of neutral buffered formalin. These samples were incubated at 4C while gently shaking for 24 hours. Immediately afterwards, samples were placed into plastic cages and processed into paraffin blocks using a Leica tissue processor with the following protocol: 70% EtOH for 1 hour x 2, 85% EtOH for 1 hour, 95% EtOH for 1 hour x 2, 100% EtOH for 1 hour x 2, Xylenes for 1 hour x 2, paraffin wax. Adipose blocks were sectioned into 6 μm sections, with 2-4 slices on each slide.

### H&E Staining

Slides were incubated at 60C for 1 hour, then placed in xylenes to remove remaining paraffin wax. Slides were then rehydrated using successive decreasing EtOH concentrations (xylenes x 2, 100% EtOH x 2, 95% EtOH, 70% EtOH, H2O). Slides were incubated in hematoxylin (Leica Surgipath 3801570), Define (3803590), Blue Buffer 8 (3802915), and eosin (3801616), and dehydrated (95% EtOH, 100% EtOH, xylene x 2). Imaging was performed using the Zeiss AxioPlan2 microscope and Olympus DP software. Analysis of adipocyte size was performed using ImageJ. Images were converted to black and white and skeletonized to reveal only the cell wall outlines. Cell area was calculated from outlines with a lower limit of 50 um and upper limit of 700 um to reduce noise. All cells from a cohort (4-7 images each from 4 animals per cohort, equal numbers of males and females) were pooled for cell area density analysis. A Welch’s unequal variances t-test was performed between ages in each diet to determine significant differences.

### Immunofluorescence

Slides were incubated at 60C for 1 hour, then placed in xylenes to remove remaining paraffin wax. Slides were then rehydrated using successive decreasing EtOH concentrations (xylenes x 2, 50% EtOH in xylenes, 100% EtOH x 2, 95% EtOH, 70% EtOH, 50% EtOH, 0.3% H2O2 in MeOH, H2O). Slides were washed with TBS and blocked in 10% normal donkey serum (Abcam ab7475) for 1 hour, followed by incubation with primary antibody overnight at 4C. [Primary antibodies: rabbit anti-Ucp1 (1:100, Sigma U6382) and mouse anti-PHH3 (1:100, Invitrogen MA5-15220)]. After an additional wash, secondary antibody was applied for 1 hour at room temperature [Secondary antibodies: donkey anti-rabbit 488 (1:1000, Abcam ab150061) and donkey anti-mouse 647 (1:200, Abcam ab150107)]. Fluoroshield Mounting Medium with DAPI (Abcam) was applied to seal the coverslip and slides were stored at 4C. Imaging was performed using the Zeiss Confocal microscope and Zen Lite imaging program. PHH3 analysis was performed using the CellProfiler program. Background was subtracted from DAPI and PHH3 channels using ImageJ. DAPI channel was used to identify total nuclei in CellProfiler. Adipose nuclei images were overlaid with PHH3 stain to identify mitotic adipose nuclei. Mitotic nuclei were summed across all 4 slides for each individual. Mitotic adipose index is reported as mitotic adipose nuclei divided by adipose nuclei multiplied by 100%.

### Quantitative rt-PCR

Total RNA was extracted from brown, subcutaneous inguinal, and visceral reproductive adipose samples using the Qiagen RNeasy Lipid Kit. High-Capacity cDNA Reverse Transcription Kit (Thermofisher) was used for reverse transcription. Quantitative-rtPCR was performed to assess expression levels of target genes with an Applied Biosystems (USA) QuantStudio 6 Flex instrument using SYBR Green reagent. Results were normalized to *L32* expression, which was experimentally determined to not be differentially expressed across diet and age cohorts. cDNA products were analyzed using the ΔCt method. Primers used: *L32* forward TCCACAATGTCAAGGAGCTG, reverse GGGATTGGTGACTCTGATGG; *Cidea* forward TGCTCTTCTGTATCGCCCAGT, reverse GCCGTGTTAAGGAATCTGCTG; *Tbx1* forward GGCAGGCAGACGAATGTTC, reverse TTGTCATCTACGGGCACAAAG; *Ucp1* forward CCTCTCCAGTGGATGTGGTAA, reverse AGAAGCCACAAACCCTTTGA.

### Mitochondrial DNA quantification

DNA was extracted from brown and inguinal adipose tissues using the Qiagen DNeasy Blood and Tissue Kit. Briefly, 40mg of tissue was homogenized in 10% proteinase K through vortexing and incubation at 56°C. DNA was precipitated with ethanol, collected in a spin column, and eluted in 150mL of buffer. DNA concentration was quantified on a Nanodrop, and 50ng was used in a qPCR reaction to quantify the amount of *h19* (nuclear gene) and *CytB* (mitochondrial gene). Mitochondrial content was calculated as the ratio of mtDNA to nucDNA. Primers used: *Cytb* forward TCTACGCTCAATCCCCAATAAAC, reverse TTAGGCTTCGTTGCTTTGAGGT; *h19* forward TATGTGCCATTCTGCTGCGA, reverse AAGGTTTAGAGAGGGGGCCT.

### RNA sequencing and analyses

Sixty-four LG/J and SM/J mice were used for sequencing analysis, representing 4 males and 4 females from each diet (high and low fat) and age (20 and 30 weeks). Total RNA was isolated from interscapular brown and reproductive white adipose tissues using the RNeasy Lipid Tissue Kit (QIAgen). RNA concentration was measured via Nanodrop and RNA quality/integrity was assessed with a BioAnalyzer (Agilent). RNAseq libraries were constructed using the RiboZero kit (Illumina) from total RNA samples with RIN scores >7.5. Libraries were checked for quality and concentration using the DNA 1000LabChip assay (Agilent) and quantitative PCR, according to manufacturer’s protocol. Libraries were sequenced at 2×100 paired end reads on an Illumina HiSeq 4000. After sequencing, reads were demultiplexed and assigned to individual samples.

FASTQ files were filtered to remove low quality reads and aligned against LG/J and SM/J custom genomes using STAR (47,48). Briefly, LG/J and SM/J indels and SNVs were leveraged to construct strain-specific genomes using the GRC38.72-mm10 reference as a template. This was done by replacing reference bases with alternative LG/J and SM/J bases using custom python scripts. Ensembl R72 annotations were adjusted for indel-induced indexing differences for both genomes. Read counts were normalized via upper quartile normalization and a minimum normalized read depth of 10 was required. Alignment summaries are provided in **Supplemental Figure 7**. Library complexity was assessed and differential expression between each age cohort for each strain-by-diet comparison was determined after TMM normalization in edgeR (49).

Functional enrichment of differentially expressed genes was tested by over-representation analysis in the WEB-based Gene Set Analysis Toolkit v2019 (50). We performed analyses of gene ontologies (biological process, cellular component, molecular function), pathway (KEGG), and phenotype (Mammalian Phenotype Ontology). For each tissue, the list of all unique differentially expressed genes was analyzed against the background of all unique genes expressed in that tissue (**Supplemental Tables 2 and 3**). A Benjamini-Hochberg FDR-corrected p-value ≤ 0.05 was considered significant.

### Correlation structure

Co-expression was assessed for the set of 62 differentially expressed cytokines and ECM genes by correlating expression of each gene with the expression of the other 61 genes in each diet-by-age cohort. Each pair of genes then had their correlations correlated (*R_g_*), where gene = *G*.

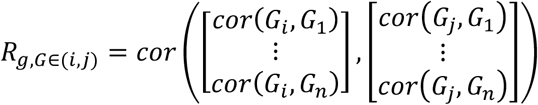

Gene-pair-correlations were then compared between the high fat-fed 30 week-old cohort and the other three cohorts (high fat-fed 30 weeks to high fat-fed 20 weeks, high fat-fed 30 weeks to low fat-fed 30 weeks, high fat-fed 30 weeks to low fat-fed 20 weeks) to obtain the Δ*R_g_* between a pair of cohorts, where cohort = K.

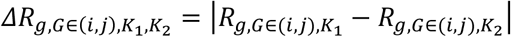

The median change in correlation (*MΔR_g_*) was calculated and permutation was employed to identify the background of expected *MΔR_g_* values. Permutation was performed by randomly selecting 2 groups of 8 animals from any cohort 10,000 times.

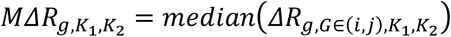

*MΔR_g_* was determined for the 2 randomized groups (*rK*_1_,*rK*_2_) for all 10,000 permutations to generate a null model. Log transformation was performed to approximate normality, which was determined by Wilks-Shapiro test and Q-Q plot. Significance was drawn from the cumulative normal null model to test if the difference in correlation structure between each pair of cohorts was greater than by chance under the randomized null model.

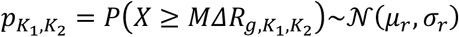

### Brown adipose excision

Interscapular brown adipose tissue depots were removed from 20 or 30 week-old high fat-fed SM/J mice. A small longitudinal incision was made between the shoulder blades. All interscapular adipose tissue was carefully removed, and a cauterizing wand used to stop excessive bleeding when necessary. Surgeries were performed under general anesthesia by IP injection of ketamine/ xylazine (100/200 mg/Kg) and mice were maintained in the surgical plane by isofluorane/oxygen for the duration of the procedure. Incisions were closed with 5-0 nonabsorbable sutures. Ketoprofen (2-5 mg/Kg) was provided post-procedure and topical antibiotic was applied to the incision for up to 3 days as necessary. Animal health and well-being was monitored daily. Sutures were removed at 10 days post-surgery. Mice were allowed to recover for four weeks after surgery or until they reached 30 weeks of age, then underwent a glucose tolerance test and an insulin tolerance test one week later. After an additional week of recovery, animals were sacrificed and plasma and multiple tissues harvested (reproductive and inguinal adipose depots, liver, heart, soleus, pancreas, hypothalamus) as described above.

### Statistics

Data within individual cohorts were assessed for normality using a Wilks-Shapiro test. Outliers were identified by a Grubbs test (p < 0.05) and removed. Data were tested for significant differences among cohorts by ANOVA with a Tukey’s post-hoc correction. The sex X diet X age term was not significant for any phenotype so males and females were pooled for analyses. P-values <0.05 were considered significant. All statistical analyses were performed using the R software package.

## Author Contributions

HAL and CC designed the experiments. CC, HS, JPW, and HAL performed physiological and molecular assays. CC, MAM and SO performed histological assays and analyses. CC and SG performed excision surgeries. CC, JFM and HAL analyzed the RNAseq data. CC and HAL wrote the manuscript. All authors edited and approved the final draft.

## List of Supplementary Figures

**Supplemental Figure 1:** Physiological parameters of the LG/J inbred mouse strain.

**Supplemental Figure 2:** Brown adipose tissue quantification of LG/J mice.

**Supplemental Figure 3:** SM/J adipose histology parameters and images

**Supplemental Figure 4:** SM/J expression of thermogenesis related genes

**Supplemental Figure 5:** SM/J thermogenic parameters

**Supplemental Figure 6:** Expression patterns of key cytokine and extracellular matrix-related genes in SM/J brown and white adipose tissue

**Supplemental Figure 7:** STAR alignment summaries for RNA-sequencing results

## List of Supplementary Tables

**Supplemental Table 1:** High and low fat diet constituents.

**Supplemental Table 2:** Differential expression results in SM/J mice.

**Supplemental Table 3:** Differential expression results in LG/J mice.

**RNA sequencing count data available for download at**: http://lawsonlab.wustl.edu/data/

## References

1. Meigs JB, Wilson PWF, Fox CS, Vasan RS, Nathan DM, Sullivan LM, et al. Body mass index, metabolic syndrome, and risk of type 2 diabetes or cardiovascular disease. J Clin Endocrinol Metab [Internet]. 2006 Aug [cited 2016 Aug 31];91(8):2906–12. Available from: http://www.ncbi.nlm.nih.gov/pubmed/16735483

2. Klöting N, Fasshauer M, Dietrich A, Kovacs P, Schön MR, Kern M, et al. Insulin-sensitive obesity. Am J Physiol Metab. 2010;

3. Goossens GH. The Metabolic Phenotype in Obesity: Fat Mass, Body Fat Distribution, and Adipose Tissue Function. Obes Facts. 2017;

4. Saito M, Okamatsu-Ogura Y, Matsushita M, Watanabe K, Yoneshiro T, Nio-Kobayashi J, et al. High incidence of metabolically active brown adipose tissue in healthy adult humans: Effects of cold exposure and adiposity. Diabetes. 2009;

5. Chondronikola M, Volpi E, Børsheim E, Porter C, Annamalai P, Enerbäck S, et al. Brown adipose tissue improves whole-body glucose homeostasis and insulin sensitivity in humans. Diabetes. 2014;

6. Gunawardana SC, Piston DW. Reversal of type 1 diabetes in mice by brown adipose tissue transplant. Diabetes. 2012;

7. Lawson H a, Zelle KM, Fawcett GL, Wang B, Pletscher LS, Maxwell TJ, et al. Genetic, epigenetic, and gene-by-diet interaction effects underlie variation in serum lipids in a LG/JxSM/J murine model. J Lipid Res. 2010;

8. Lawson HA, Lee A, Fawcett GL, Wang B, Pletscher LS, Maxwell TJ, et al. The importance of context to the genetic architecture of diabetes-related traits is revealed in a genome-wide scan of a LG/J × SM/J murine model. Mamm Genome [Internet]. 2011 Apr [cited 2016 Jan 20];22(3-4):197–208. Available from: http://www.pubmedcentral.nih.gov/articlerender.fcgi?artid=3650899&tool=pmcentrez&rendertype=abstract

9. Ehrich TH, Kenney JP, Vaughn TT, Pletscher LS, Cheverud JM. Diet, obesity, and hyperglycemia in LG/J and SM/J mice. Obes Res. 2003;

10. Ahren W and. The High-Fat Diet–Fed Mouse. Diabetes. 2004;

11. Shimizu I, Aprahamian T, Kikuchi R, Shimizu A, Papanicolaou KN, MacLauchlan S, et al. Vascular rarefaction mediates whitening of brown fat in obesity. J Clin Invest. 2014;

12. Shimizu I, Walsh K. The Whitening of Brown Fat and Its Implications for Weight Management in Obesity. Current obesity reports. 2015.

13. Roberts-Toler C, O’Neill BT, Cypess AM. Diet-induced obesity causes insulin resistance in mouse brown adipose tissue. Obesity. 2015;

14. Lapa C, Arias-Loza P, Hayakawa N, Wakabayashi H, Werner RA, Chen X, et al. Whitening and Impaired Glucose Utilization of Brown Adipose Tissue in a Rat Model of Type 2 Diabetes Mellitus. Sci Rep. 2017;

15. Wu J, Boström P, Sparks LM, Ye L, Choi JH, Giang AH, et al. Beige adipocytes are a distinct type of thermogenic fat cell in mouse and human. Cell. 2012;

16. Kim D, Kim J, Yoon JH, Ghim J, Yea K, Song P, et al. CXCL12 secreted from adipose tissue recruits macrophages and induces insulin resistance in mice. Diabetologia. 2014;

17. Mejhert N, Wilfling F, Esteve D, Galitzky J, Pellegrinelli V, Kolditz CI, et al. Semaphorin 3C is a novel adipokine linked to extracellular matrix composition. Diabetologia. 2013;

18. Sun K, Tordjman J, Clément K, Scherer PE. Fibrosis and adipose tissue dysfunction. Cell Metabolism. 2013.

19. Lee JT, Pamir N, Liu NC, Kirk EA, Averill MM, Becker L, et al. Macrophage metalloelastase (MMP12) regulates adipose tissue expansion, insulin sensitivity, and expression of inducible nitric oxide synthase. Endocrinology. 2014;

20. Huang G, Ge G, Wang D, Gopalakrishnan B, Butz DH, Colman RJ, et al. α3(V) Collagen is critical for glucose homeostasis in mice due to effects in pancreatic islets and peripheral tissues. J Clin Invest. 2011;

21. Edén D, Siegbahn A, Mokhtari D. Tissue factor/factor Viia signalling promotes cytokine-induced beta cell death and impairs glucose-stimulated insulin secretion from human pancreatic islets. Diabetologia. 2015;

22. Gauger KJ, Bassa LM, Henchey EM, Wyman J, Bentley B, Brown M, et al. Mice deficient in Sfrp1 exhibit increased adiposity, dysregulated glucose metabolism, and enhanced macrophage infiltration. PLoS One. 2013;

23. D’souza AM, Neumann UH, Glavas MM, Kieffer TJ. The glucoregulatory actions of leptin. Molecular Metabolism. 2017.

24. Alcalá M, Calderon-Dominguez M, Bustos E, Ramos P, Casals N, Serra D, et al. Increased inflammation, oxidative stress and mitochondrial respiration in brown adipose tissue from obese mice. Sci Rep. 2017;

25. Abdullah A, Peeters A, de Courten M, Stoelwinder J. The magnitude of association between overweight and obesity and the risk of diabetes: A meta-analysis of prospective cohort studies. Diabetes Res Clin Pract. 2010;

26. Strazzullo P, D’Elia L, Cairella G, Garbagnati F, Cappuccio FP, Scalfi L. Excess body weight and incidence of stroke: Meta-analysis of prospective studies with 2 million participants. Stroke. 2010.

27. Kenchaiah S, Evans JC, Levy D, Wilson PWF, Benjamin EJ, Larson MG, et al. Obesity and the Risk of Heart Failure. N Engl J Med. 2002;

28. Rauscher GH, Mayne ST, Janerich DT. Relation between body mass index and lung cancer risk in men and women never and former smokers. Am J Epidemiol. 2000;

29. Reeves GK, Pirie K, Beral V, Green J, Spencer E, Bull D. Cancer incidence and mortality in relation to body mass index in the Million Women Study: Cohort study. Br Med J. 2007;

30. Centers for Disease Control and Prevention. National Diabetes Statistics Report. US Dep Heal Hum Serv. 2017;

31. Yang W, Dall TM, Beronjia K, Lin J, Semilla AP, Chakrabarti R, et al. Economic costs of diabetes in the U.S. in 2017. Diabetes Care. 2018;

32. Colditz GA, Willett WC, Rotnitzky A, Manson JE. Weight gain as a risk factor for clinical diabetes mellitus in women. Ann Intern Med. 1995;

33. Chatterjee S, Khunti K, Davies MJ. Type 2 diabetes. The Lancet. 2017.

34. Dulloo AG, Montani JP. Pathways from dieting to weight regain, to obesity and to the metabolic syndrome: An overview. Obesity Reviews. 2015;

35. Tomiyama AJ, Ahlstrom B, Mann T. Long-term Effects of Dieting: Is Weight Loss Related to Health? Soc Personal Psychol Compass. 2013;

36. Nathan DM, Barrett-Connor E, Crandall JP, Edelstein SL, Goldberg RB, Horton ES, et al. Longterm effects of lifestyle intervention or metformin on diabetes development and microvascular complications over 15-year follow-up: The Diabetes Prevention Program Outcomes Study. Lancet Diabetes Endocrinol. 2015;

37. MacArthur JW. Genetics of Body Size and Related Characters. I. Selecting Small and Large Races of the Laboratory Mouse. Am Nat. 2002;

38. Poekes L, Lanthier N, Leclercq IA. Brown adipose tissue: a potential target in the fight against obesity and the metabolic syndrome. Clin Sci. 2015;

39. Kajimura S, Spiegelman BM, Seale P. Cell Metabolism Review Brown and Beige Fat: Physiological Roles beyond Heat Generation. Cell Metab. 2015;

40. Moreno-Viedma V, Amor M, Sarabi A, Bilban M, Staffler G, Zeyda M, et al. Common dysregulated pathways in obese adipose tissue and atherosclerosis. Cardiovasc Diabetol. 2016;

41. Hasegawa Y, Ikeda K, Chen Y, Alba DL, Stifler D, Shinoda K, et al. Repression of Adipose Tissue Fibrosis through a PRDM16-GTF2IRD1 Complex Improves Systemic Glucose Homeostasis. Cell Metab. 2018;

42. Maquoi E, Munaut C, Colige A, Collen D, Roger Lijnen H. Modulation of adipose tissue expression of murine matrix metalloproteinases and their tissue inhibitors with obesity. Diabetes. 2002;

43. Raz V, Sterrenburg E, Routledge S, Venema A, van der Sluijs BM, Trollet C, et al. Nuclear entrapment and extracellular depletion of PCOLCE is associated with muscle degeneration in oculopharyngeal muscular dystrophy. BMC Neurol. 2013;

44. Shu L, Chan KHK, Zhang G, Huan T, Kurt Z, Zhao Y, et al. Shared genetic regulatory networks for cardiovascular disease and type 2 diabetes in multiple populations of diverse ethnicities in the United States. PLoS Genet. 2017;

45. Takahashi N, Yoshizaki T, Hiranaka N, Kumano O, Suzuki T, Akanuma M, et al. The production of coagulation factor VII by adipocytes is enhanced by tumor necrosis factor-α or isoproterenol. Int J Obes. 2015;

46. Denroche HC, Kwon MM, Glavas MM, Tudurí E, Philippe M, Quong WL, et al. The role of autonomic efferents and uncoupling protein 1 in the glucose-lowering effect of leptin therapy. Mol Metab. 2016;

47. Nikolskiy I, Conrad DF, Chun S, Fay JC, Cheverud JM, Lawson HA. Using whole-genome sequences of the LG/J and SM/J inbred mouse strains to prioritize quantitative trait genes and nucleotides. BMC Genomics [Internet]. 2015 Jan [cited 2016 Jan 20];16:415. Available from: http://www.pubmedcentral.nih.gov/articlerender.fcgi?artid=4445795&tool=pmcentrez&rendertype=abstract

48. Dobin A, Davis CA, Schlesinger F, Drenkow J, Zaleski C, Jha S, et al. STAR: ultrafast universal RNA-seq aligner. Bioinformatics [Internet]. 2013 Jan 1 [cited 2016 Jul 27];29(1):15–21. Available from: http://www.ncbi.nlm.nih.gov/pubmed/23104886

49. Chen Y, Mccarthy D, Robinson M, Smyth GK. edgeR: differential expression analysis of digital gene expression data User’s Guide. 2015.

50. Zhang B, Kirov S, Snoddy J. WebGestalt: An integrated system for exploring gene sets in various biological contexts. Nucleic Acids Res. 2005;

