## Supplemental Figures for "Brown adipose expansion and remission of glycemic dysfunction in obese SM/J mice"

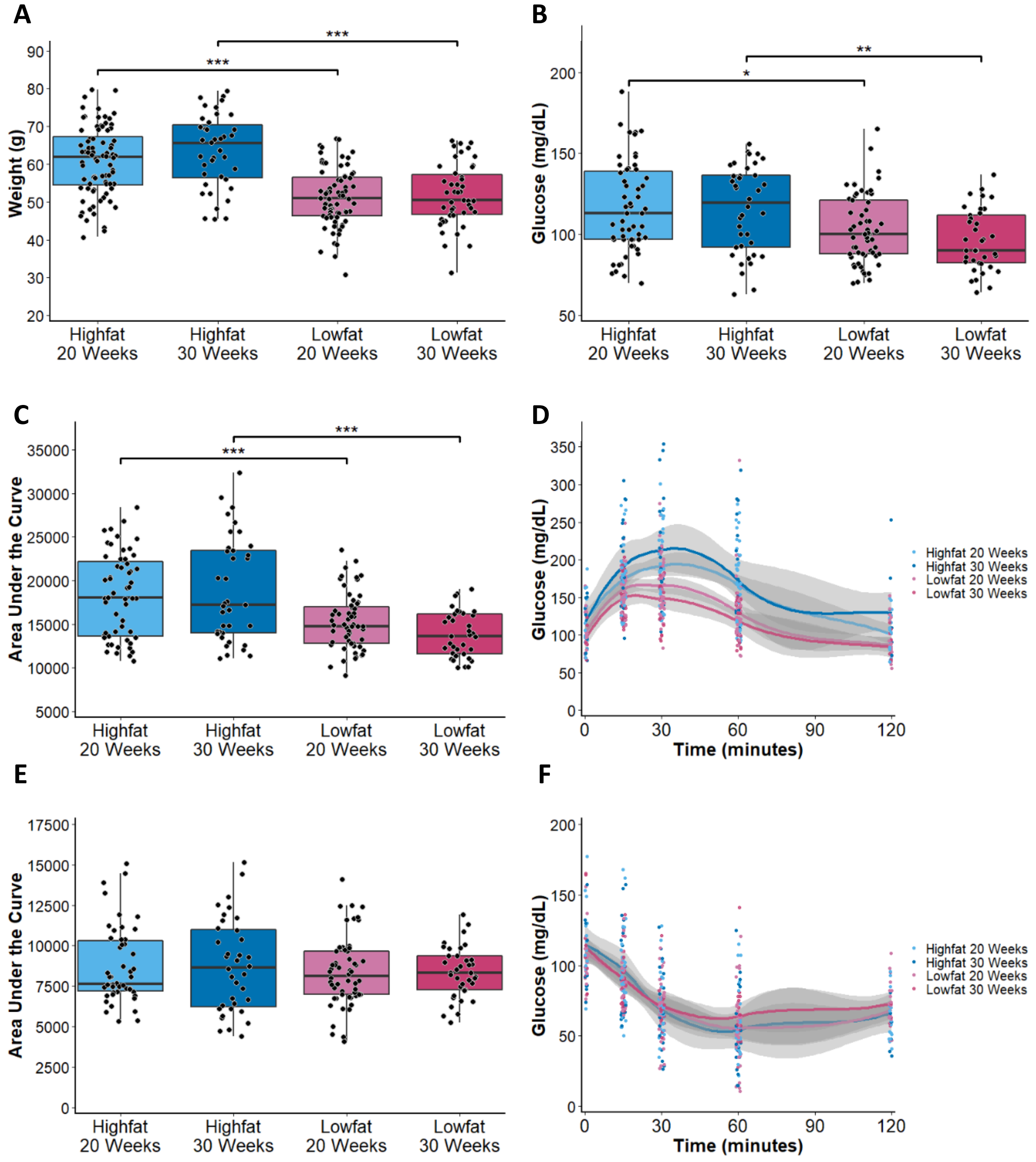


**Supplemental Figure 1: Physiological parameters of the LG/J inbred mouse strain.**

**A** Body weight of high and low fat-fed LG/J mice at 20 and 30 weeks of age. **B** Basal glucose and (**C, D)** glucose tolerance of high fat-fed LG/J mice are higher than low fat-fed controls at 30 weeks. **E-F** Insulin sensitivity is not different among the LG/J cohorts. Panels A-F n = 30-60. Equal numbers of males and females represented. All measurements taken identically to SM/J mice as described in main Methods. *p<0.05, **p<0.01, ***p<0.0001

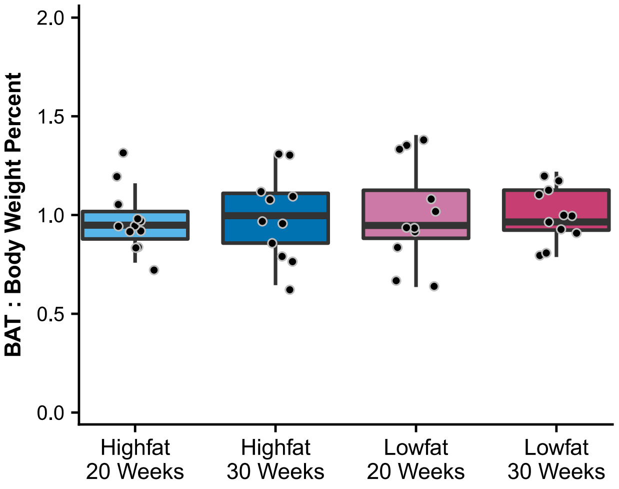


**Supplemental Figure 2: Brown adipose tissue quantification of LG/J mice.**

Quantification of intrascapular brown adipose depot weight reported as percentage of total weight of LG/J mice on high and low fat diets at 20 and 30 weeks of age. n = 11-12 mice, equal numbers of males and females. Brown adipose tissue and body weight determined using the same methods as for SM/J mice described in main Methods section.

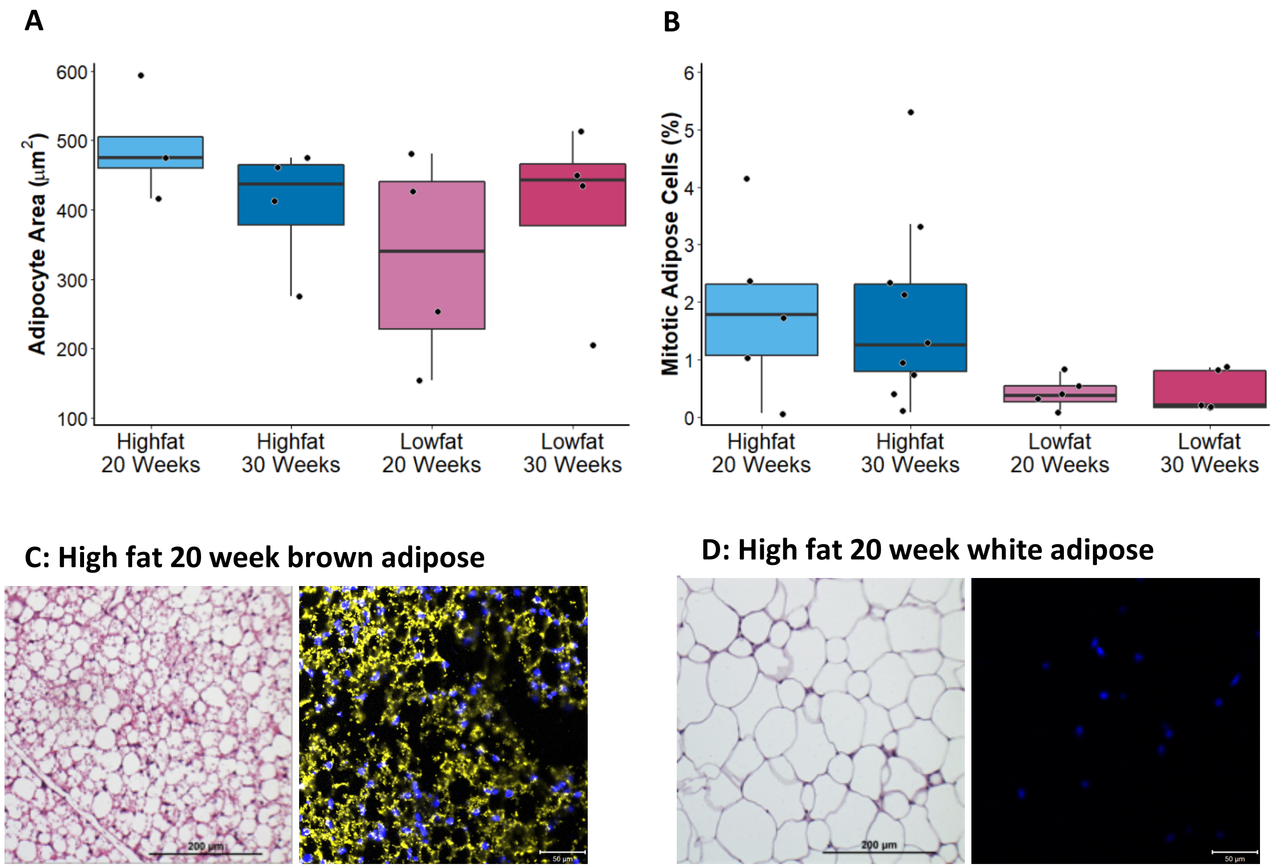

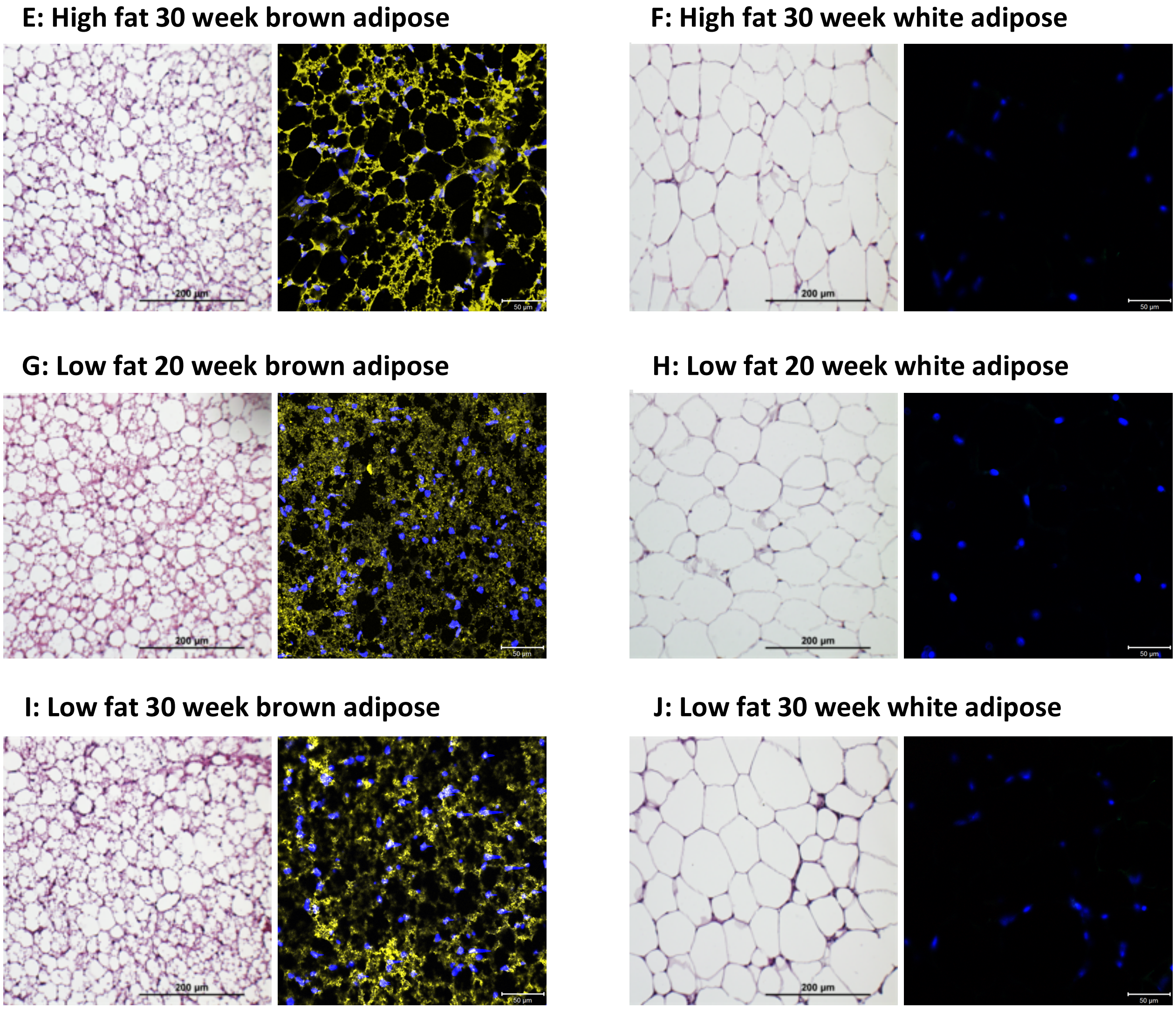


**Supplemental Figure 3: SM/J adipose histology**

Brown adipose sections from high and low fat-fed mice at 20 and 30 weeks were stained with hematoxylin and eosin (H&E) (**A**) or DAPI to mark nuclei and a phosphohistone H3 (pHH3) antibody to mark mitotic nuclei (**B**). Four images were taken per animal. ImageJ was used to determine average cell area per animal. Cellprofiler was used to count total number of nuclei and number of pHH3-positive nuclei. Total number of pHH3-positive nuclei was summed for each individual animal and represented as a percentage of total number of nuclei. Representative images of brown (left panels) and white (right panels) adipose from (**C-D**) high fat-fed mice at 20 and (**E-F**) 30 weeks of age and from (**G-H**) low fat-fed mice at 20 and (**I-J**) 30 weeks of age. Stained for H&E (left picture) or UCP1 (yellow) and DAPI (blue) (right picture). Histology procedures described in main Methods section.

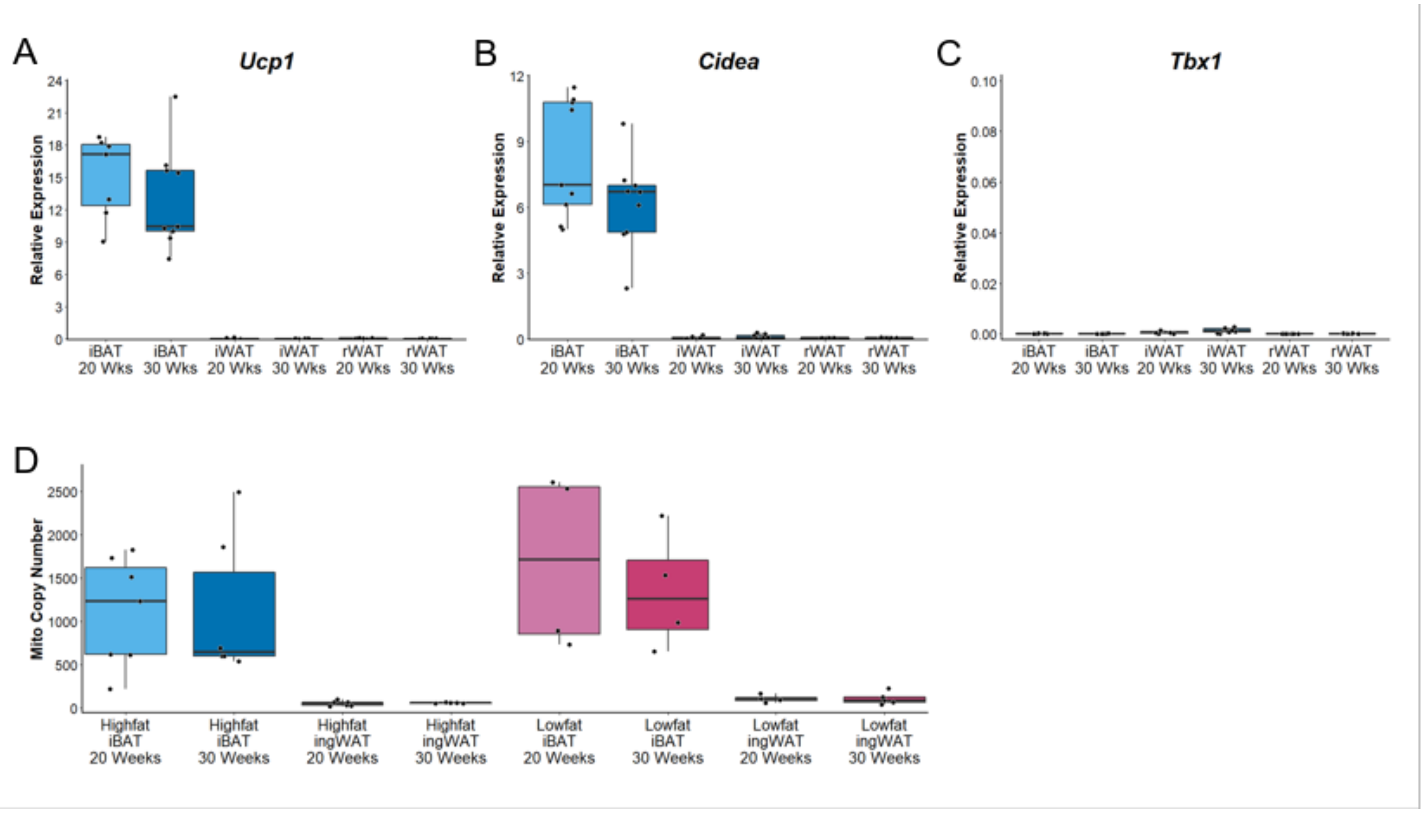


**Supplemental Figure 4: Thermogenic genes do not change between 20 and 30 week adipose tissue in high fat SM/J animals.**

Gene expression levels quantified in three adipose depots of high fat-fed mice: intrascapular brown adipose (iBAT), inguinal white adipose (ingWAT), and reproductive white adipose (repWAT), n = 6-10 mice per cohort and tissue. Canonical brown adipose genes (**A**) Ucp1 and (**B**) Cidea show high expression in iBAT and no difference between 20 and 30 week-old mice. Beige adipose marker (**C**) Tbx1 is not expressed in any depot. (**D**) Mitochondrial copy number was significantly higher in brown adipose tissue than in inguinal white adipose tissue at both 20 and 30 week time points with no significant difference between high or low fat-fed mice, n = 6-7 mice per cohort and tissue. Equal numbers of males and females represented. * $p<0.05$, ** $p< 0.01$

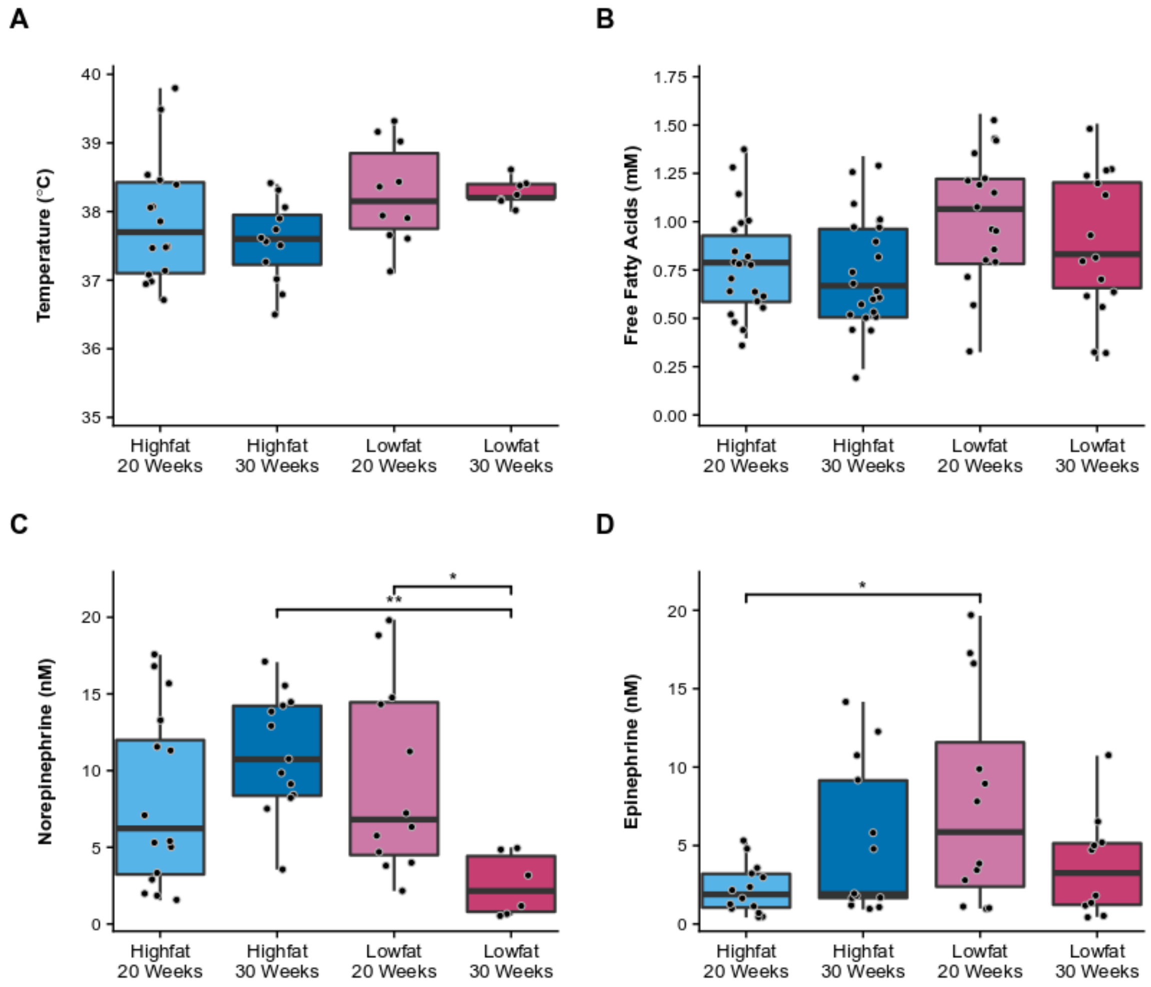


**Supplemental Figure 5: SM/J thermogenic parameters**

**A** Body temperature and (**B**) plasma free fatty acid concentration are not different among SM/J cohorts. **C** Plasma norepinephrine and (**D**) epinephrine do not change between 20 and 30 weeks in high fat-fed mice. Body temperature was taken at time of necropsy using a rectal probe calibrated to room temperature (OakTon Temp JKT, Acorn). Plasma free fatty acids, norepinephrine, and epinephrine were analyzed from blood collected at necropsy. For free fatty acid assay, plasma was collected and quantified according to main methods section using the Wako HR Series NEFA-HR Color Reagent A (NC9517308) and Color Reagent B (NC9517310). For the catecholamine assays, 150 µL of blood was prepared according to protocol from Vanderbilt University Medical Center's Hormone Assay and Analytical Services Core (www.vumc.org/hormone/assays). Panels A, C, D n = 8-16, panel B n = 15-21mice, equal numbers of males and females represented. $^*p<0.05$, $^{**}p<0.01$, $^{***}p<0.0001$

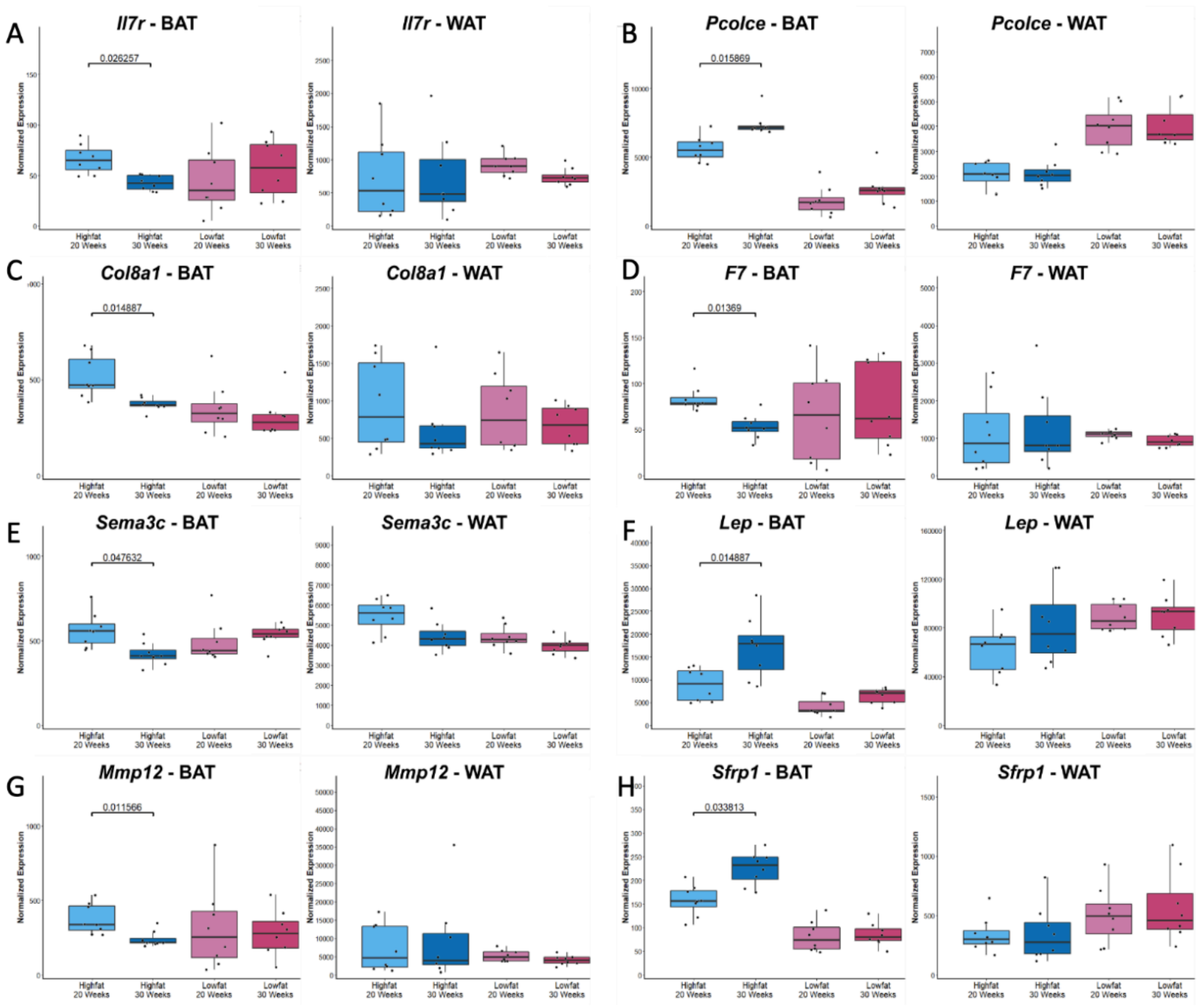


**Supplemental Figure 6: Expression patterns of key cytokine and extracellular matrix-related genes in SM/J brown and white adipose tissue.**

Normalized expression for 8 key cytokine and extracellular matrix related genes (A = *Il7r*, B = *Pcolce*, C = *Col8a1c*, D = *F7,* E = *Sema3*, F = *Lep*, G = *Mmp12*, H = *Sfrp1*) in SM/J brown (left) and white (right) adipose tissue. Fdr-corrected p-values are provided for differential expression by age within the SM/J high fat cohort. BAT = brown adipose tissue; WAT = white adipose tissue

**A**

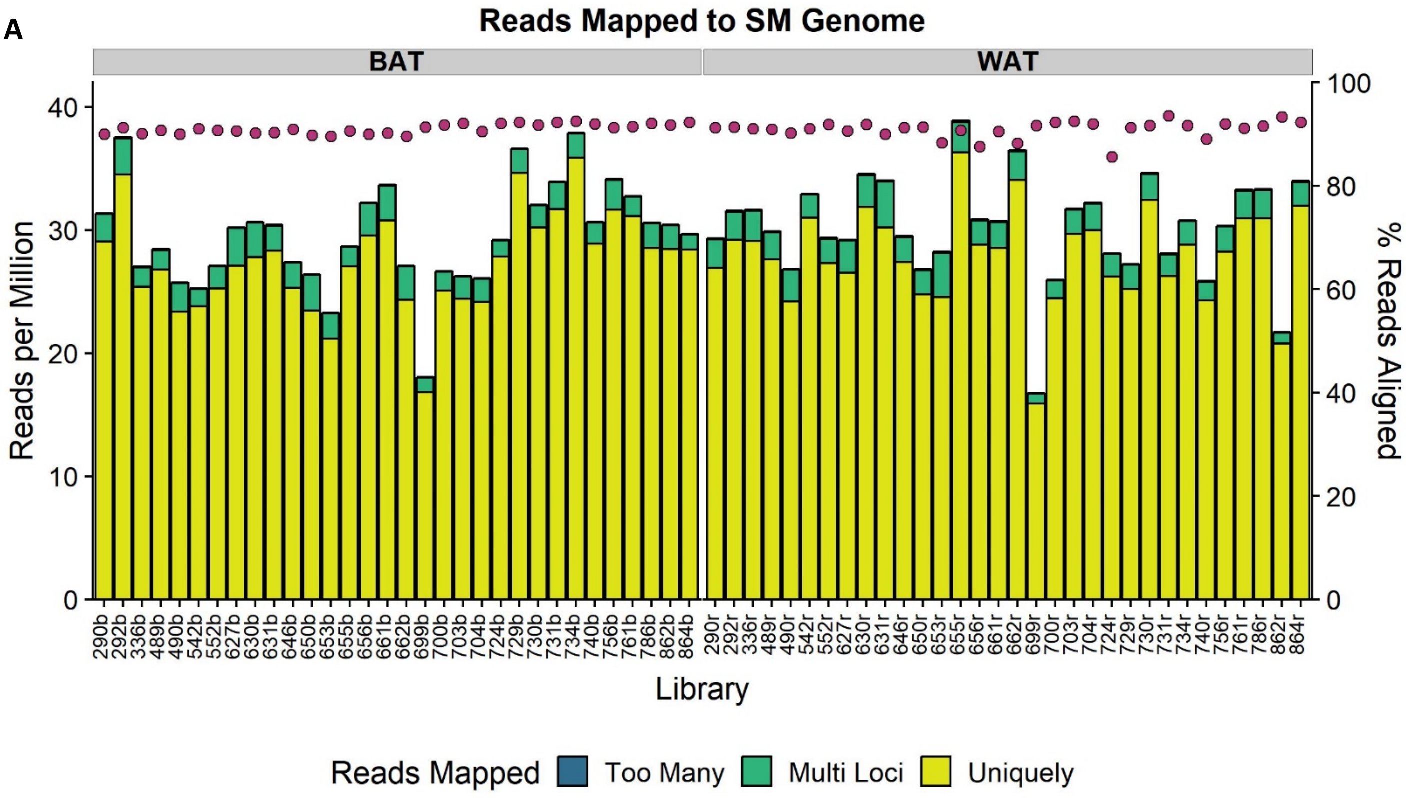


**B**

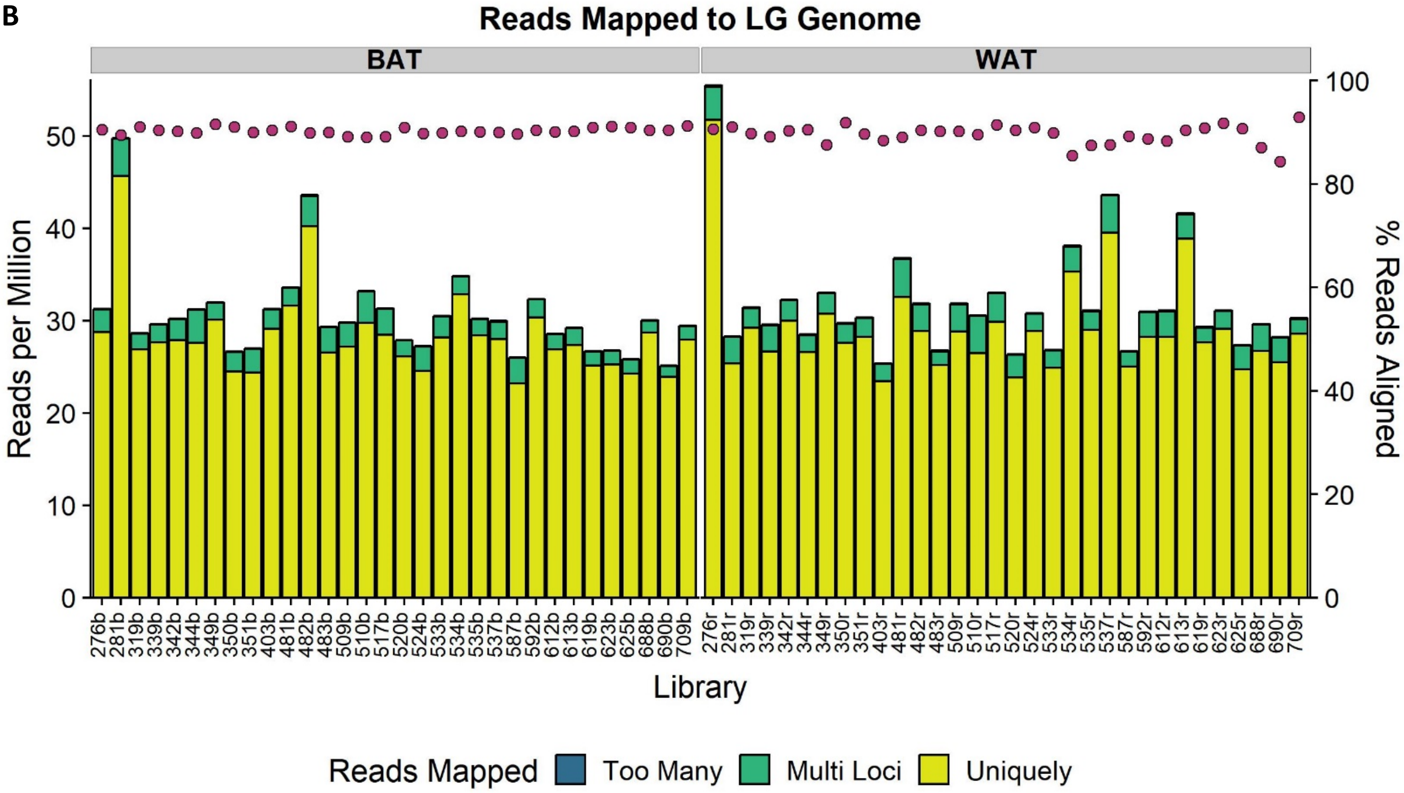


**Supplemental Figure 7: STAR alignment summaries for RNA-sequencing results**

Alignment summaries for RNA-sequencing results from (**A**) SM/J and (**B**) LG/J samples. Brown adipose samples are on the left, white adipose are on the right. Reads per million (left y-axis) are indicated by bars (yellow = uniquely mapped, green = multi loci, blue = mapped to too many loci) and % reads aligned (right x-axis) are indicated by red circles.

**Supplemental Table 1: High and low fat diet constituents.** The diets (high fat: Teklad TD88137; low fat: Research Diets D12284) have been used in multiple studies of diet-induced obesity using the SM/J strain.

**Supplemental Table 2: Differential expression results in SM/J mice.** Includes Ensembl ID, log fold change (logFC) between 20 and 30 weeks, raw p-value (PValue) and FDR-corrected p-value (FDR) for differential expression, and average normalized read counts for 20 (Avg20) and 30 (Avg30) weeks. Tabs for high fat brown adipose (SM_HF_BAT), high fat white adipose (SM_HF_WAT), low fat brown adipose (SM_LF_BAT), and low fat white adipose (SM_LF_WAT).

**Supplemental Table 3: Differential expression results in LG/J mice.** Includes Ensembl ID, log fold change (logFC) between 20 and 30 weeks, raw p-value (PValue) and FDR-corrected p-value (FDR) for differential expression, and average normalized read counts for 20 (Avg20) and 30 (Avg30) weeks. Tabs for high fat brown adipose (LG_HF_BAT), high fat white adipose (LG_HF_WAT), low fat brown adipose (LG_LF_BAT), and low fat white adipose (LG_LF_WAT).
