## Supplemental Table 1 for "Brown adipose expansion and remission of glycemic dysfunction in obese SM/J mice"

| **Supplemental Table 1. High and low fat diet constituents** | | |
| --- | --- | --- |
| Source | High Fat^#^ | Low Fat^*^ |
| Energy from Fat | 42% | 15% |
| Casein (g/kg) | 195 | 197 |
| Sugars (g/kg) | 341 | 307 |
| Corn starch (g/kg) | 150 | 313 |
| Cellulose (g/kg) | 50 | 30 |
| Corn oil (g/kg) | --- | 58 |
| Hydrogenated coconut oil (g/kg) | --- | 7 |
| Anhydrous milkfat (g/kg) | 210 | --- |
| Cholesterol (g/kg) | 1.5 | --- |
| Total energy (kJ/g) | 18.95 | 16.99 |
| ^#^ Teklad TD88137  ^*^ Research Diets D12284 | | |
